# MEA-seqX: High-resolution Profiling of Large-scale Electrophysiological and Transcriptional Network Dynamics

**DOI:** 10.1101/2024.05.15.594367

**Authors:** Brett Addison Emery, Xin Hu, Diana Klütsch, Shahrukh Khanzada, Ludvig Larsson, Ionut Dumitru, Jonas Frisén, Joakim Lundeberg, Gerd Kempermann, Hayder Amin

**Author notes:** shared first authorship (Brett Addison Emery, Xin Hu). shared senior authorship (Gerd Kempermann, Hayder Amin).

## Abstract

Concepts of brain function imply congruence and mutual causal influence between molecular events and neuronal activity. Decoding entangled information from concurrent molecular and electrophysiological network events demands innovative methodology bridging scales and modalities. Our MEA-seqX platform, integrating high-density microelectrode arrays, spatial transcriptomics, optical imaging, and advanced computational strategies, enables the simultaneous recording and analysis of molecular and electrical network activities at the level of individual cells. Applied to a mouse hippocampal model of experience-dependent plasticity, MEA-seqX unveiled massively enhanced nested dynamics between transcription and function. Graph-theoretic analysis revealed an increase in densely connected bimodal hubs, marking the first observation of coordinated spatiotemporal dynamics in hippocampal circuitry at both molecular and functional levels. This platform also identified different cell types based on their distinct bimodal profiles. Machine-learning algorithms accurately predicted network-wide electrophysiological features from spatial gene expression, demonstrating a previously inaccessible convergence across modalities, time, and scales.

## Introduction

The brain evolved to process complex information robustly and efficiently to maintain homeostasis, navigate the world, make decisions, and perform higher cognitive functions^1^. Understanding the complexity of the brain coherently from the molecular to system level requires integrating multimodal data from diverse spatiotemporal contexts^1^. At the core of this ambitious objective lies the integration of neuronal electrophysiological and molecular phenotypes at the network and cellular resolution, as these underlie physiological functions as well as neurodevelopmental and neurodegenerative diseases^2,3^. Towards this aim, methodological developments like patch-sequencing (patch-seq) have allowed single-cell transcriptomics and the morphologic reconstruction of individual neurons after electrophysiological recordings^4,5^. Despite its extraordinary significance, patch-seq still suffers from very low throughput and shortcomings in resolving neuronal networks in medium to large spatial contexts. The Electro-seq method combined flexible bioelectronics with *in situ* RNA sequencing to map electrical activity and gene expression but is only applicable to reductionist neuronal cultures^6^.

Two independent developments, however, allow a new perspective on this problem. By profiling expression patterns of thousands of neuronal genes while preserving spatial tissue architecture, high-throughput spatially resolved transcriptomics (SRT) offers unparalleled insights into the molecular diversity of brain regions of interest^7,8^. However, the temporal resolution of SRT is limited, providing only single snapshots of the transcriptomic landscape. Spatiotemporal transcriptomics is only resolvable using multiple samples from different time points^9,10^. The functional state to which the molecular signature relates has to be inferred from circumstantial evidence. This clearly falls short of appreciating the diversity of electrical and molecular signals and their interplay underlying brain function, cognition, and behavior^11,12^. But novel brain-on-chip technologies empowered by high-density complementary metal-oxide semiconductors (CMOS) biosensing microelectrode arrays (CMOS-MEA) are now allowing the non-invasive, multi-site, long-term, and label-free simultaneous measurements of extracellular activity capturing both local field potentials and spiking activity from thousands of neurons with single-cell accuracy without disruption of cellular integrity^13–17^. Merging these cutting-edge technologies would potentiate the insight to be gained from decoding the spatiotemporal electrophysiological and transcriptomics information in the same tissue while retaining cell positioning. The underlying fundamental hypothesis is that brain functions are executed through the joint action of large assemblies of neurons and gene networks that share basic organizational principles^18^ and information processing across a wide range of spatial and temporal scales^19,20^ that evolve with experience and change in disease.

To test this prediction, we present here MEA-seqX, which combines brain-on-chip, bioimaging, and spatial sequencing technologies within a cross-scale computational framework. MEA-seqX allows simultaneous recording of electrophysiological firing patterns from large cell assemblies in acute brain slices at high spatiotemporal resolution, imaging the entire circuit for spatial localization, and multiplexed profiling of the cellular transcriptomics from the same neural circuit. Using automatic machine-learning algorithms and preserving time and topology in one high-resolution representation, MEA-seqX can specify spatiotemporal transcriptomic networks in their real-time relation to connectivity and other multimodal data, quantify molecular dynamics in pseudotime based on the underlying firing information, deconvolute spatially resolved cell type compositions, and predict electrophysiological network features from transcriptomic profiles with high accuracy.

To illustrate the power of this approach, we apply it to the classical enriched environment paradigm of experience-dependent plasticity, in which the sole experimental intervention lies in differential housing conditions of laboratory mice. The enriched mice live in a larger group of isogenic animals in a larger enclosure compared to standard-housed animals^16,21^. This straightforward yet hugely influential paradigm elicits structural and functional changes throughout the brain and hippocampus, shaping scientific and public discourse. We have recently demonstrated that its effects on an unexpected scale relate to changes in the hippocampal circuit level^16^. MEA-seqX has now enabled us to explore a once inaccessible question - how the computational dynamics and connectome of a large-scale hippocampal network are connected to the underlying transcriptional dynamics. Our hypothesis posited a causal link between these two dynamics, a connection frequently implied in biomedical contexts but previously supported by only sparse and limited data points. MEA-seqX changes this by providing a more robust foundation.

Our study unveils the potential of identifying the molecular identity and dynamics of large-scale neural circuits. We envision the development of new multimodal models of high functional validity in the contexts of health and disease. Such “biomarkers” hold immense potential for the development of novel diagnostic and screening tools, especially in but not limited to precision medicine.

## Results

### Interfacing Technologies and Integrating Information – from Transcriptome to Functional Networks

MEA-seqX integrates brain-on-chip technology via network electrophysiology (n-Ephys) of high-density CMOS-microelectrode array (CMOS-MEA)^13,16^, bioimaging via optical microscopy, and spatial sequencing technology via spatially resolved transcriptomics (SRT) (**Figure 1a-d**). Specifically, high-resolution spatiotemporal recordings of extracellular firing patterns obtained from 300 µm mouse hippocampal-cortical ‘HC’ acute slices interfaced with 4096-on-chip sensors. Simultaneously, optical imaging was employed for precise anatomical localization, and high-resolution transcriptomic profiling data was obtained from the same cells within the circuit.

**Figure 1.**
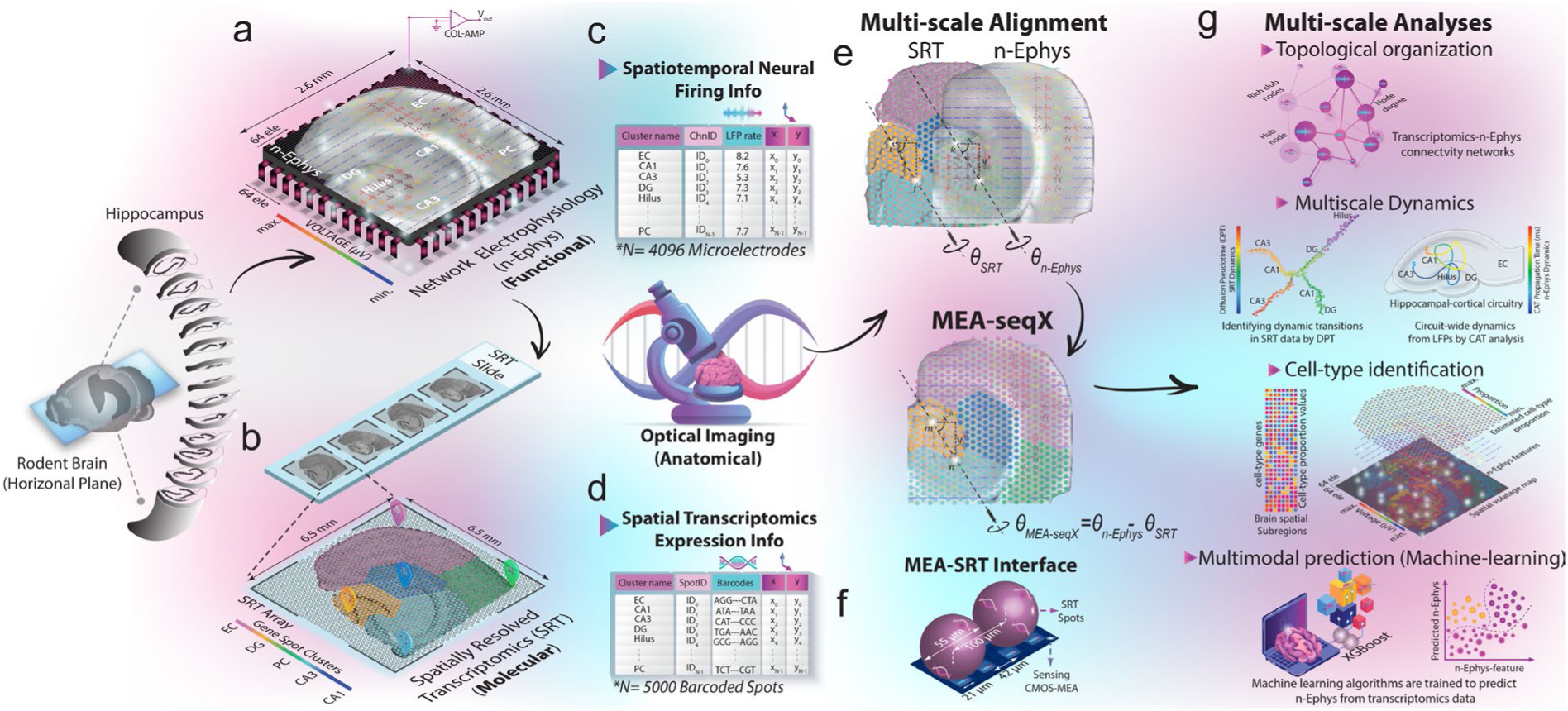
MEA-seqX Platform Overview for Integrating Brain Dynamics Across Scales. **a, b)** The platform combines high-density CMOS-MEA, optical microscopy, and spatial sequencing technology to examine the relationship between spatial transcriptomics and neural oscillatory dynamics. **c)** A high-density CMOS-MEA equipped with 4096 sensing electrodes adeptly captures functional firing LFP patterns stemming from all HC regions with pinpoint precision in spatial coordinates. **d)** Leveraging spatial barcodes, SRT with 5000 spots allows the systematic mapping of gene expression profiles, elucidating the spatial nuances across HC regions. **e)** A Python-based pipeline processes data to map transcriptional-functional dynamics. This includes a multifaceted spatial rescaling and alignment approach, establishing a direct association between neural activity and transcriptomic features. **f)** A closer examination, exemplified by a sensing electrode coupled with the SRT spot, highlights the discernible pitch size difference between the two platforms. **g)** Advanced analysis from multidimensional readouts provides insights into multiscale network features. The analysis, including topological graphs capturing the intricate web of multimodal transcriptional-functional connectivity. Also, multiscale neural dynamics are quantified with gene pseudotime and center of activity trajectories. Cell-type composition is inferred from spatial transcriptomic data correlated with their firing characteristics obtained from n-Ephys. Additionally, the automatic machine-learning algorithm accurately predicts electrophysiological features from transcriptomic profiles.

Quality control of our transcriptomic datasets revealed similar nFeature and nCount RNA statistics and tissue structure when visualized via a Uniform Manifold Approximation and Projection (UMAP) method^22^ (**Supplementary Figures 1a-c**). Network-wide activity in the HC was assessed through Principal Component Analysis (PCA) and K-means clustering algorithms^13^, which provided distinct features of oscillatory waveforms and their shapes in each interconnected HC layer (**Supplementary Figure 1d**). The multidimensional readouts were processed through a Python-based computational pipeline to quantitatively map the molecular dynamics of the circuit at high spatiotemporal resolution. A multiscale spatial rescaling and alignment procedure was developed to establish a direct correspondence between the n-Ephys electrode-SRT spot interface and their respective network-wide functional electrical activity and transcriptomic feature readouts with their localization in the tissue. This was implemented with automatic scaling algorithms, employing image resizing and rotation based on optical imaging, n-Ephys electrode-SRT spot interface physical size, and two anatomical landmarks in the dentate gyrus (**Figure 1e**; for details, see Methods). The resulting overlay allowed the identification of network features from the transcriptomics readouts in alignment with readouts of neural activity (**Figure 1g**). MEA-seqX generated precise topological graphs of multimodal data connectivity, presenting both the local and global relationships of SRT spots and the underlying firing electrodes (**Figure 1g**). From the wealth of high-dimensional data, transcriptional pseudotime dynamics underlying firing information flow were derived (**Figure 1g**). By integrating a deconvolution method, we inferred cell-type resolution from the spatial transcriptomic data and correlated the neuronal heterogeneity to their firing features (**Figure 1g**). Finally, MEA-seqX provided an automatic machine-learning algorithm to predict high-accuracy network electrophysiological features from spatial transcriptomic profiles (**Figure 1g**).

### Linking Spatial Transcriptome to Network-wide Neural Functional Dynamics

To apply the pipeline to a concrete, previously unsolvable research question, we used the MEA-seqX framework to uncover the impact of experience-dependent plasticity^16^ on the coordinated activity of neuronal ensembles and their interaction with orchestrated transcriptional activity. HC slices from mice housed in standard (SD) and enriched environment (ENR) conditions were prepared for recording oscillatory patterns of local field potentials (LFPs), optical imaging, and SRT sequencing. To assess how spatial patterns of gene expression (SRT) correspond to the functional network electrophysiological features (n-Ephys) of the same brain tissue, we measured Spearman’s correlation^23^ to quantify the transcriptional similarity of gene expression profiles from SRT spots and to examine their relationship to functional network features (i.e., amplitude, delay, energy, LFP rate, negative peak count, and positive peak count). We found a significant enhancement in the gene expression pattern corresponding to the functionally coupled dentate gyrus (DG) and CA3 regions in ENR compared to SD (**Figure 2a**). In particular, the LFP rate showed a 2.2-fold, 2.5-fold, and 0.6-fold increase of correlated transcripts in the DG, CA3, and CA1 subfields, respectively, in the ENR compared to the SD (**Figure 2b**). We next sought to identify which specific genes would drive a stronger causal link between correlated molecular and functional networks. Based on gene ontology clusters, we categorized correlated genes into six targeted gene families, including Immediate Early Genes (IEGs), Hippocampal Neurogenesis, Hippocampal Signaling Pathway, Receptors and Channels, Synaptic Plasticity, Synaptic Vesicles, and Adhesion^24,25^. Examining transcripts from these families illustrated enhanced expression of IEGs, ion channel activity, synaptic function, and neurogenesis in ENR compared to SD. Genes essential for hippocampal activity and function, such as *Bdnf*, *Egr1*, *Homer1, Npas4, Gria2*, and *Campk2a*, had higher expression levels in the hippocampal transcriptome of ENR compared to SD (**Supplementary Figure 2**). Next, to identify distinct spatiotemporal patterns across the combined SRT and n-Ephys modalities, we implemented an unsupervised machine-learning algorithm using a sparsity-constrained non-negative matrix factorization (NMF)^26^ This approach allowed decomposing modalities into sets of differentially expressed subnetworks of genes, spatial locations, and electrophysiological features to provide dimensionality reduction and interpretability (**Figure 2c**). IEGs exhibited a significant contribution to linking spatial transcription patterns to LFP activity (**Supplementary Figure 2**). To determine the optimal number of components (i.e., factors *p*) to be discerned by the dimensionality reduction model within the NMF decomposition, we evaluated the efficacy of reconstructing V-matrix across a spectrum of model complexities. A distinct “elbow” was identified as situated between two linear regimes of the reconstruction error^27^ at = 12. At this point, incorporating additional patterns into the model resulted in marginal enhancements in the fit quality (**Figure 2d**). The input expression V-matrix, which contains the combined information from SRT and n-Ephys (i.e., LFP rate) data, with each entry representing the expression level of a gene correlated to network function feature at a specific spatial location (**Figure 2e**), was decomposed into two non-negative matrices. The basis W-matrix contains the spatial gene expression patterns and their locations rated to the functional feature (**Figure 2f**), and the coefficient H-matrix represents the contribution of these basis vectors to each spatial location and n-Ephys feature (**Figure 2g**). This analysis unveiled increased normalized gene expression and network feature values in the ENR network, along with higher spatially resolved components driving the strength in gene expression. Additionally, we found spatially specific subnetworks in the SD and ENR networks, highlighting genes such as *Bdnf*, *Egr1*, *Fosb*, and *Npas4*.

**Figure 2.**
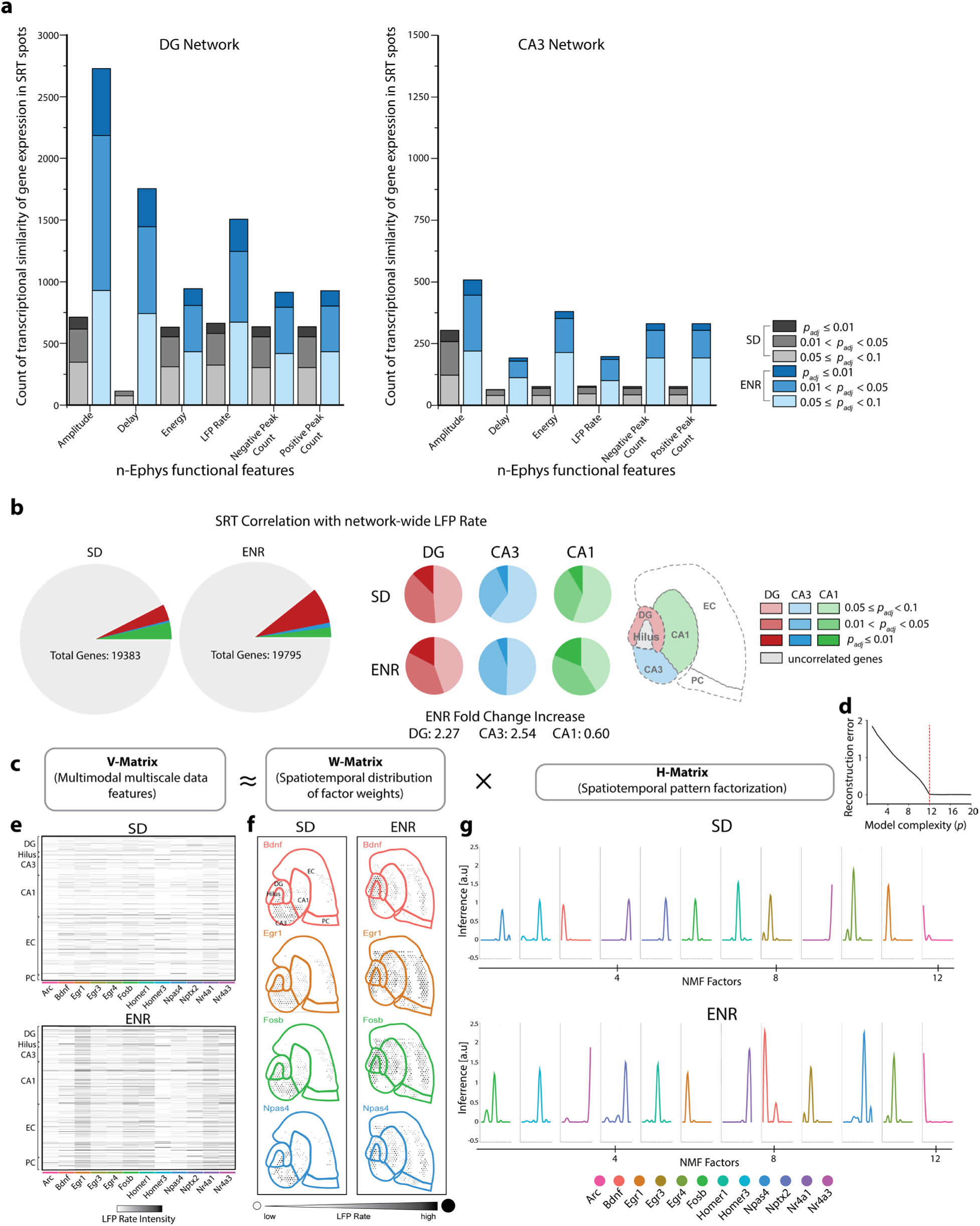
Integration of Spatial Gene Expression Patterns and Network Electrophysiological Features. **a)** Comparative analysis of SRT expression patterns and functional n-Ephys features illustrate enhanced gene expression patterns in ENR corresponding to the DG and CA3 regions compared to SD. The statistical significance of SRT and n-Ephys features correlation is quantified using Benjamini-Hochberg false-discovery rate (FDR)-adjusted *p-values* (*padj*). **b)** Quantitative profiling of correlated genes based on network-wide LFP rate reveals a 2.2-fold increase in correlated genes in the DG and 2.5-fold and 0.6-fold increases in CA3 and CA1 subregions, respectively, in ENR compared to SD. The significance of SRT and LFP correlation is measured using Benjamini-Hochberg FDR-adjusted *p-values* (*padj*). **c)** Application of unsupervised machine learning through sparsity-constrained NMF. The algorithm enables the joint analysis of SRT and n-Ephys data, revealing diverse subnetworks of genes, spatial locations, and electrophysiological features. **d)** The reconstruction error of V, considering various quantities of shared spatiotemporal patterns in H. An evident “elbow” is observed at n = 12, where the enhancement of the model diminishes. **e)** The combined input expression matrix (V) captures the collective information from SRT and n-Ephys (LFP rate) datasets. Each entry signifies the gene expression level correlated to network function features at a specific spatial location. **f)** Visualization of spatial gene expression patterns through the basis matrix (W). This matrix illustrates how gene expression patterns are distributed across spatial locations and linked to functional features. **g)** Coefficient matrix (H) representing the contribution of basis vectors to spatial locations and n-Ephys features. The analysis reveals increased normalized gene expression and network feature values in the ENR network, emphasizing spatially resolved components driving gene intensity.

MEA-seqX suggests a computational role of the experience-dependent dynamics in the coordinated interaction of neuronal ensemble activity and their corresponding transcription patterns.

### Coordinated Topological Network Organization of Spatially-resolved Transcriptome and Activity Patterns

We employed quantitative measures to comprehensively examine the interconnections between HC subnetworks derived from SRT data and neural n-Ephys recordings under SD and ENR conditions. We calculated ‘mutual information’^28^, i.e., the measure of the mutual dependence between two variables, in order to assess the extent of the interdependence of gene expression within specific gene families (**Supplementary Figure 3a**) and Pearson’s correlation coefficient (PCC)^13,16^ to gauge the cross-covariance among pairs of firing electrodes (**Supplementary Figure 3b**). These computations allowed us to establish connectivity matrices on multiple scales. Next, we computed mutual information distance scores for each target gene family in every spot to gauge the dissimilarities in interaction within the different spots. These scores were then compared between spots and organized into clusters^28^ (**Supplementary Figures 4a-f**). Simultaneously, we quantified differences in the correlation matrices by analyzing the PCC of concurrent LFP activity across interconnected HC layers^13,16^ (**Supplementary Figure 4g**). Upon analyzing various gene families, it became evident that the ENR transcriptome, in contrast to SD, exhibited higher mutual information. This suggests a more robust statistical relationship in coordinated activity and communication among gene expression patterns within the HC subnetworks (**Supplementary Figures 4a-f**). Importantly, as evident from the LFP cross-correlogram^16^ (**Supplementary Figure 4g**), this finding strongly aligned with the significant enhancement in both the local and global strength of spatiotemporal interactions in ENR vs. SD networks^16^.

Subsequently, in order to describe and quantify the topological organization, flow of information, and communication properties within the multimodal transcriptional-functional readouts, MEA-seqX employed a graph-theoretic approach to assessing multiscale network metrics (**Figure 3a**). This involved constructing detailed wiring diagrams depicting both local and global interconnections between transcriptomics and neural activity patterns. The transcriptomic graph was formed using mutual information scores, while the functional graph was generated from the connectivity patterns among co-firing neuronal ensembles, as captured by the LFPs under SD and ENR conditions. The resultant connectivity maps between transcriptomics and neural function revealed the spatial arrangement of interconnected subnetworks derived from spatial IEGs and spatiotemporal functional neural activity. Remarkably, these maps demonstrated analogous spatial distributions and connectivity patterns across different scales and modes (**Figures 3b-c**). Comparing HC networks between SD and ENR further emphasized a heightened level of across-scale causal coordination. The transcriptional-functional connectome was enhanced in ENR compared to SD.

**Figure 3.**
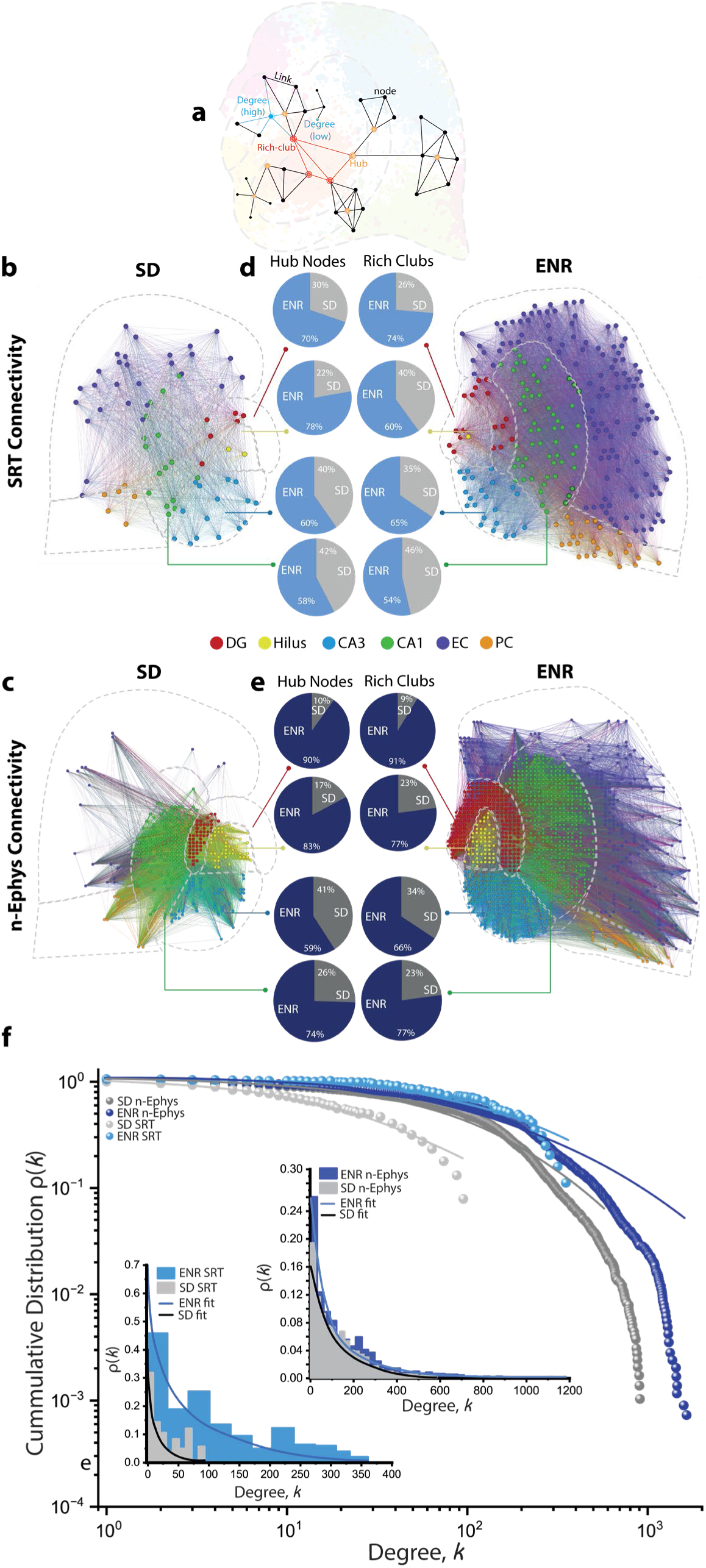
Multiscale Network Topological Analysis of Spatially Sequenced Connectome and Neural Functional Connectivity. **a)** Illustration of key multiscale graph measures depict the characteristic Hub and Rich-club nodes defined based on their degree of interconnectedness in an acute hippocampal-cortical slice. Node degree corresponds to the number of attached links at a given node. **b)** Connectivity maps of spatial IEGs in HC-interconnected layers in ENR and SD. The networks are visualized with Gephi to illustrate total connections from complete sequenced SD data (node = 546, and links = 5166) and ENR data (node = 877, and links = 50974). **c)** Connectivity maps of spatiotemporal functional neural activity in HC-interconnected layers under ENR and SD conditions. The networks are visualized with Gephi to illustrate 2% of total connection in SD (node = 1057, and links = 24217) and ENR large-scale recorded data (node = 2003, and links = 36312). Graph nodes in (b) and (c) are scaled according to degree strength and colored according to HC module association and indicated in colored circles legends. Colored links identify the intra- and inter-cluster connections. **d)** The percent of quantified hubs and rich-club nodes in different hippocampal transcriptomic networks in SD and ENR. **e)** The percent of quantified hubs and rich-club nodes in different hippocampal functional networks in SD and ENR. **f)** The power-law distributions indicate scale-free transcriptomic (SRT)-functional (n-Ephys) network topologies with small-world properties in SD and ENR networks. The log-log plot of the cumulative connection distribution for ENR (SRT and n-Ephys; *blue*) networks exhibits a significantly heavier tail than SD networks (SRT and n-Ephys; *grey*), indicating low-degree nodes coexist with a few densely connected hubs, yet higher than SD networks, which reach a cut-*Kolmogorov-Smirnov test*). This is also supported at the linear scale (insets) for all conditions, and their compliance to power-law is assessed with Pareto fits (*\*p< 0.05 Kolmogorov-Smirnov test*). The lognormal function fitted the power-law distributions with goodness-of-fit in a log-log plot (with a coefficient of determination *R^2^* = 0.95, 0.98, 0.96, and 0.97 for SRT, n-Ephys (SD), SRT, n-Ephys (ENR), respectively). The probability density function of Pareto fitted the power-law distributions with goodness-of-fit in the linear plots (with *R^2^* = 0.96, 0.97, 0.95, and 0.97 for SRT, n-Ephys (SD), SRT, n-Ephys (ENR), respectively).

We also conducted analyses on the constructed graphs to identify highly interconnected nodes termed “hub complexes” and densely connected hubs referred to as “rich-club organization” within the transcriptional-functional connectomes^29^. ENR subnetworks, derived from both SRT and n-Ephys data, exhibited greater interconnected hub complexes and rich-club organization (**Figures 3d, e; *blue***) compared to their SD counterparts (**Figures 3d, e; *grey***). This indicates that enriched experience leads to enhanced specialization in coordination interactions, heightened resilience, and an increased capacity for global communication^30^ across transcriptional-functional scales. These results shed new light on the dynamic interactions and mutual influences between molecular and functional hub complexes, contributing significantly to the overall coordination of multiscale topological network organization.

Many biological networks are characterized by a small-world topology^31^ defined by a scale-free architecture consisting of highly connected hub nodes and a degree distribution that decays with a power-law tail^32^. By analyzing cumulative degree distributions of interconnected links in the transcriptomics and functional connectome from SD and ENR networks, we found that these multiscale distributions indeed followed a power-law function, which we have previously also postulated for a transcriptomic network of adult hippocampal neurogenesis^33^ and network-wide activity in ENR^16^. Both SRT and n-Ephys distributions in ENR networks displayed a heavier tail than SD, indicating a more significant number of densely linked hubs (**Figure 3f and insets**). This finding is supported by sparsity-constrained NMF analysis of multiscale degree distributions (i.e., similar to those implemented in Figure 2c-g). We quantified multiscale decomposed sets of variably expressed subnetworks of genes, spatial locations, and the network’s connectivity node degree features to identify the variation in network topological metrics concomitant with the expression of spatially-resolved IEGs and the degree of network connectivity (**Supplementary Figure 5**).

By revealing experience-induced hippocampal connectomics across scales and its intricate multilayer dynamic characteristics within a single experiment, our results demonstrate the capacity of MEA-seqX to integratively capture coordinated transcriptomics and functional data. This integration allows novel insights into neural communication, resilience levels, hierarchical organization, and specialization across multiple scales that previously could only be studied independently and usually based on limited data^30,34,35^.

### Assessing Multiscale Dynamics of Multimodal Information

To address the challenge of unraveling the synchronous dynamics across scales and modalities, we combined two cutting-edge computational methods – Diffusion Pseudotime (DPT)^36^ for SRT and Center of Activity Trajectories (CAT)^13,16^ for n-Ephys. This integrative approach aimed at unraveling the temporal progression of gene expression and network-wide neural activity in hippocampal circuitry. We applied DPT to the static snapshot SRT data to achieve pseudotemporal ordering by assembling the spots according to expression similarity. This allowed us to construct a network representation of SRT developmental trajectories. The probability of differentiation was computed through Euclidian distances from vector-based randomized distances in the diffusion map space, which facilitated the identification of low-dimensional changes from high-dimensional observations. We here focused on IEGs expression in SD and ENR, revealing significant regional differences in DPT based on IEGs expression (**Figures 4a, b**). Concurrently, we quantified the intra-hippocampal spatiotemporal propagation pathway by constructing CATs for all n-Ephys circuit-wide oscillatory activity and thereby calculated the rate of spatiotemporal displacement of those firing patterns (**Figures 4c, d**). The analysis of CATs duration demonstrated faster propagation (i.e., shorter duration) of firing events in ENR compared to SD, as previously reported^16^. Remarkably, the comparison of SD and ENR transcriptomes indicated a faster DPT in all four hippocampal regions of ENR transcriptome compared to SD as well, thus mirroring the faster spatiotemporal propagation patterns observed in the ENR obtained from n-Ephys CATs^16^ (**Figures c-e**). By integrating the DPT spatial maps with the temporal progression of neural activity trajectories, modulated by the impact of rich experience, allowed us to instantiate a multiscale perspective of how molecular and electrical processes unfold simultaneously and interact within a biological system. Diving deeper into experience-dependent activity dynamics, the comparison between SD and ENR becomes particularly enlightening. Such a comparison potentially unravels the causal link between dynamical changes at the molecular level and those on functional scales. Different experiential environments, as represented by SD and ENR, induce distinct responses at the transcriptome level, which are reciprocally and indivisibly linked to the neural activity in the same cells. These differential activity patterns connect transcriptomics to the function and back^37,38^.

**Figure 4.**
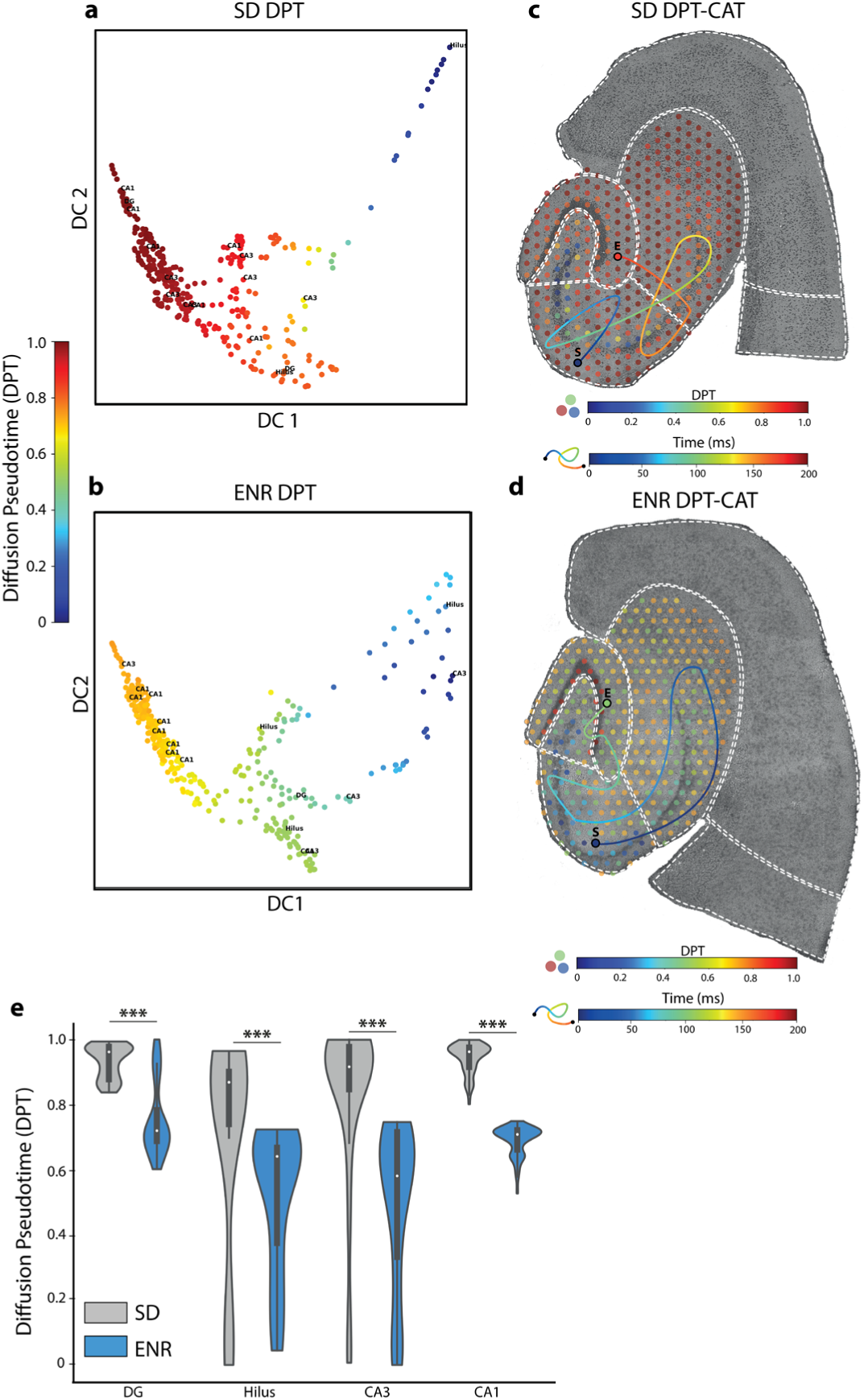
Analysis of Multiscale Hippocampal Dynamics. **a)** Differential progression of DPT in the expression of IEGs from SRT SD data showcases the developmental trajectory of cells within the hippocampal spatial regions. **b)** Same as in (a) but in ENR. **c)** Correspondence between DPT and CAT analyses in SD to infer colocalization, spatial-temporal alignment, and functional insights. **d)** Same as in (c) but in ENR. **e)** Quantifying differential DPT progression in interconnected hippocampal regions in SD and ENR (*\*p< 0.001 ANOVA test*).

### Spatiotemporal Cell-type Identification

Next, to understand the transcriptional diversity of neural cell types and their roles in the firing patterns of the hippocampal circuits^39^, we employed the conditional autoregressive-based deconvolution (CARD) method using a single-cell sequencing reference^40,41^. This allowed us to determine cell types and local tissue composition from the deconvolved gene expression patterns in order to construct multiscale spatial maps of the heterogeneity of neural types and their firing characteristics in the same hippocampal-cortical tissue. The initial application of CARD yielded a broad group classification of hippocampal cell types. This diverse group encompassed astrocytes, endothelial cells, ependymal cells, macrophages, microglial cells, neurogenic cells, neurons, oligodendrocytes, and NG2 cells. Prior to any filtering, we identified 85 different cell types, a finding that underscored the experimental validity of our slice acquisition technique. Following the removal of low-count cell types, we were left with a robust group of 76 cell types.

Interestingly, when these cell types were exposed to two different transcriptomic inputs from SD and ENR, we observed a consistent distribution of prominent cell types across both transcriptomes (**Figure 5a**). This result underscores the robustness of our approach and the reproducibility of our findings. Further, integrating the CARD method into MEA-seqX proved instrumental in achieving spatially resolved cell-type composition and linking it to large-scale oscillatory electrophysiological features (**Figure 5b**). We identified specific high-proportionate cell types within the DG and CA3 regions based on their unique marker genes. In the DG, granule cells (GCs) were characterized by *Cck* and *Penk* expressions, while CA3 pyramidal cells displayed *Nos1* and *Inhba* markers. In accordance with prior studies^42^, we have observed the presence of both common *Cck*-expressing and less frequent *Penk*-expressing GCs within the molecular layers of the DG and suprapyramidal blade (**Figure 5c**). Notably, the expression of both marker genes was significantly higher in ENR compared to SD (**Figure 5e**). When these markers were superimposed onto the DG-functional network data, the ENR samples showed enhanced firing patterns and signal amplitude, especially within the DG’s suprapyramidal blade (**Figures 5c, e**). Prior research has associated *Penk* with enrichment in DG engram cells and its involvement in hippocampal-associated behaviors^42^, while *Cck is* implicated in the dynamic selection and control of cell assemblies in DG^43^. Our data align with existing reports and highlight a marked increase in DG excitability and temporal dynamics in ENR, suggesting a potential avenue for exploring how enriched experiences might affect the transcriptional-functional interplay in hippocampal circuitry, aiding in understanding the cellular and molecular basis of memory^44^. Conducting a similar analysis, focusing on the spatial distribution of the pyramidal cell layer in CA3 (**Figure 5d**), we identified *Nos1* and *Inhba* marker genes exhibiting significantly higher expression in ENR than SD (**Figure 5f**). This transcriptomic readout matches the increased LFP rate and distinctive waveform characteristics identified within the CA3 region (**Figures 5d, f**). *Nos1* has established associations with pivotal neural mechanisms, encompassing long-term potentiation (LTP), synaptic plasticity, and regulating neural circuit dynamics^45,46^, while *Inhba* is linked to neuroprotection and neuronal survival^47^. These reports provide robust substantiation for our enhanced transcriptional-functional findings within the framework of the experience-dependent paradigm evident in the ENR group. This, in turn, paves the way for a more profound exploration of the specific role played by these marker genes within CA3 pyramidal neurons, along with its potential implications for comprehending neural function and dysregulation across scales.

**Figure 5.**
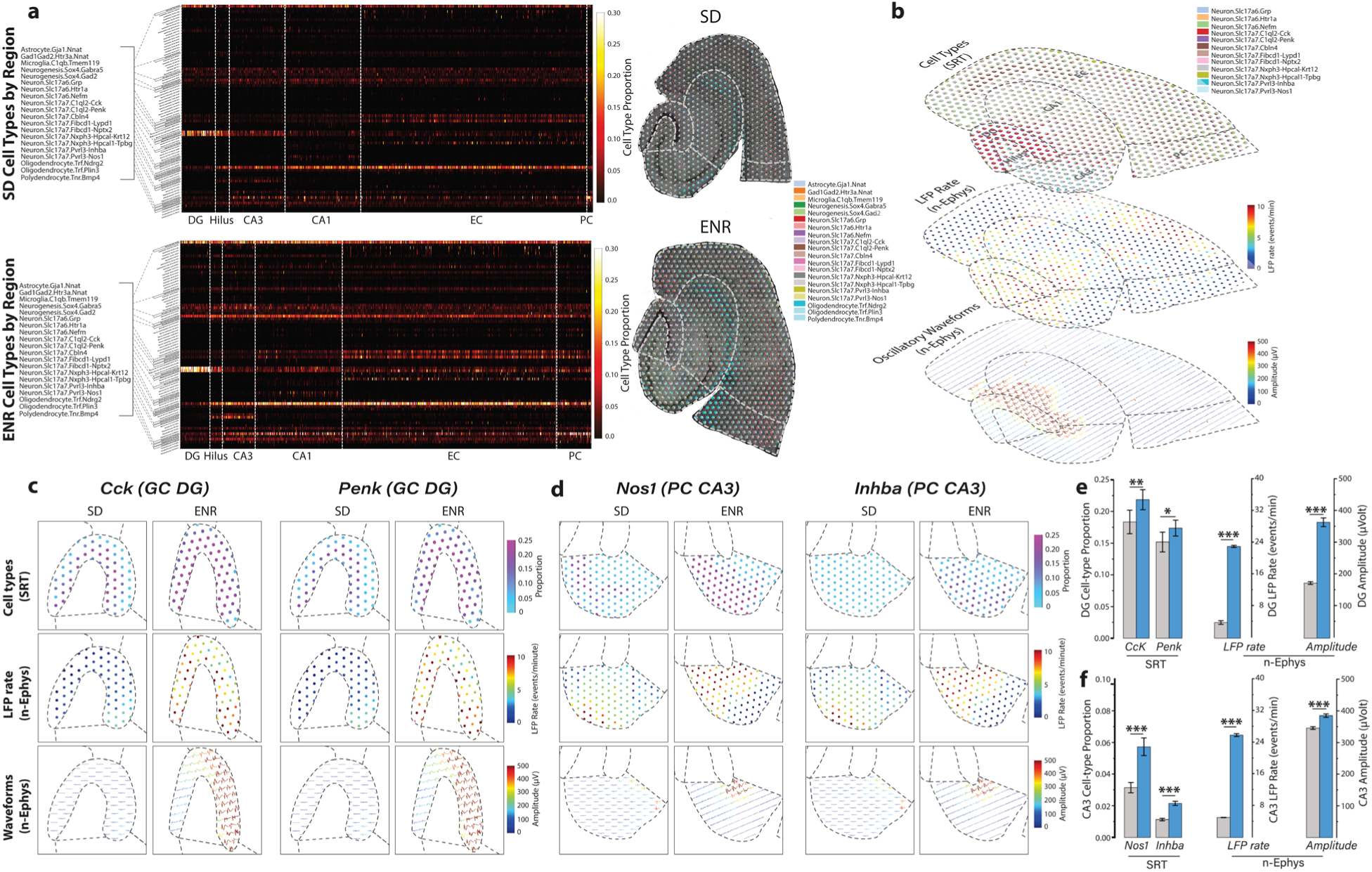
Analysis of Spatial Cell-Type and their Firing Pattern Fingerprints. **a)** A regionally ordered heatmap based on the CARD method depicts the cell type proportions of the 76 filtered cell types. Inset lists and localized cell types superimposed on SD and ENR brain slices highlight the 20 cell types with the highest proportions. Comparative analysis of hippocampal transcriptomes between SD and ENR conditions reveals a comparable distribution of prominent cell types. **b)** Integrating the CARD methodology into the MEA-seqX platform yields a spatially resolved composition of cell types accompanied by underlying oscillatory firing characteristics. This presentation includes the top 20 cell types with the highest proportions, as determined by SRT readouts, LFP Rate, and oscillatory waveforms derived from n-Ephys readouts across the entire hippocampal network. **c)** Examination focusing on the DG region offers an in-depth view of the spatially resolved composition of cell types alongside the corresponding oscillatory firing features. **d)** Similarly, a regional assessment within the CA3 pyramidal cell (PC) network provides insight into the spatial composition of cell types and their corresponding oscillatory firing features. **e)** The ENR transcriptome exhibits higher proportions of spatially localized marker genes - *Cck* and *Penk*-associated with granule cell types in the DG. These proportions are notably elevated compared to the SD transcriptome. Similarly, ENR exhibits a higher LFP rate and amplitude than SD (*\*\*\*p < 0.001*, *\*\*p < 0.01*, *ANOVA, **p < 0.05*, *ANOVA*). **f)** Analysis in the CA3 region reveals increased proportions of spatially localized marker genes *Nos1* and *Inhba*, linked to pyramidal cell types, in the ENR transcriptome compared to the SD transcriptome, which correlates with the higher LFP rate and amplitude on the functional network scale *(***p < 0.001, ANOVA*).

Moving beyond single-cell methods and focusing on multiscale network-level dynamics, our findings provide a more comprehensive and nuanced understanding of hippocampal-cortical cell types and their interactive sequential electrophysiological properties across multiple scales.

### Prediction Across Scales and Modalities

To investigate whether the expression profile of individual spatially-resolved genes can predict HC network-wide electrophysiological features, we employed the Gradient Boosting (XGBoost) Algorithm, known for its strong interpretability by integrating multiple tree models^48^. While previous methods have focused on gene properties correlated with electrophysiological and morphological diversity across cell types using low-resolution transcriptomics and electrophysiology (e.g., single-cell RNA sequencing and patch-clamp)^4,5,39^, our study aimed at assessing whether specific electrophysiological features could be predicted using spatially-resolved transcriptomic data. The XGBoost model was trained for each quantitative n-Ephys measure using 70% of spatial transcriptomic data points in the detected SRT spots (i.e., 333 genes across the spatial context of HC tissue and six gene families as input). Three spatiotemporal n-Ephys metrics (LFP rate, amplitude, and time delay) were successfully predicted based on the differential spatial gene expression. The relationship between cross-validated predictions and the ground truth was evaluated with the Pearson correlation coefficient (*r*) for SRT-spot-n-Ephys electrodes (**Figure 6a** and **Supplementary Figure 6**). By implementing the XGBoost on specific gene families that exhibited higher expression in ENR compared to SD circuits, we observed significantly higher prediction accuracy for the ENR dataset. The XGBoost classifier achieved ~93% accuracy for the ENR dataset and ~70% for the SD dataset (**Figure 6b**). Such predictive interplay between individual genes or gene families at the transcriptomic level and network-level functionality might support the idea that brain functions are orchestrated via multiscale networks that follow fundamental organizational tenets^18^. These results underscore a multiscale causative association between neural activity, plasticity, and distinct spatial gene expressions within specific gene families. This aligns with the predictions of network-level functionality modulated by prior experience^16^. This approach could shed light on critical genes and cellular pathways shaping neuronal responses and overall brain function while revealing the regulatory mechanisms governing neural dynamics, plasticity, and disease pathogenesis. Our integrated approach will massively reduce experimental complexity, thus enhancing result interpretability and offering guidance for subsequent investigations. Specific genes predicted to influence particular electrophysiological features may be targeted for manipulation to validate their functional role^49^.

**Figure 6.**
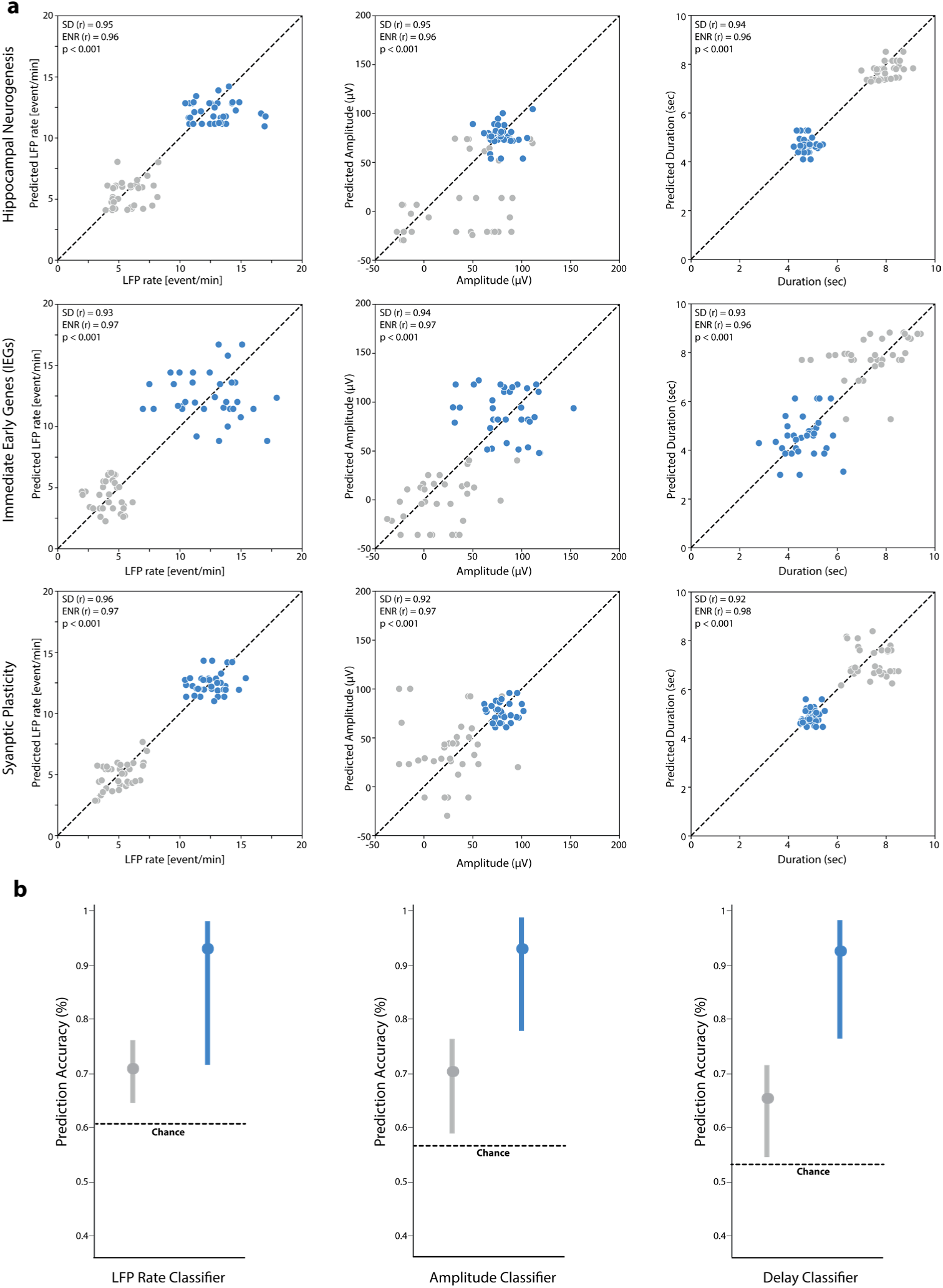
Machine-learning Prediction for Multiscale Transcriptional-Functional Data. **a)** Application of the XGBoost algorithm to predict network electrophysiological metrics (LFP rate, amplitude, and duration) from transcriptomic data of each specified gene family in SD and ENR data. The prediction of n-Ephys metrics from the transcriptomics data is evaluated with Pearson correlation coefficient (*r*) in SD and ENR. The significant difference between the predicted SD and ENR values in all gene families is indicated (*\*\*\*p < 0.001*, *ANOVA*). **b)** Performance of the XGBoost indicated by mean accuracy value comparison from final data output iterations of all gene families exhibiting higher prediction accuracy for the ENR than SD data. The values are computed over the mean, and three standard deviations are determined to be within the threshold of chance.

### Conclusions and Outlook

In this study, we introduced the MEA-seqX platform, which provides the unprecedented capability to simultaneously capture and integrate molecular and functional information across multiple spatiotemporal scales within intact brain tissues. MEA-seqX pushes the boundaries beyond existing technologies such as patch-seq, electro-seq, and spatial transcriptomics by combining the advantages of various techniques while overcoming their individual limitations. MEA-seqX presents a distinctive solution to limited spatiotemporal resolution by harnessing the high-density capabilities of CMOS-MEA technology combined with spatial transcriptomics, optical imaging, and computational tools, offering exceptional temporal resolution down to individual cells in their spatial contexts. The platform fills a crucial gap in our understanding of the molecular infrastructure supporting and resolving the integrity of large-scale neuronal interactions in physiological and experience-dependent plasticity paradigms^18,50^. Through our integrative approach, we identified spatially resolved causal regulation across molecular-functional features, which uncovered the impact of environmental factors on coordinated neural activity and gene expression – a suspected but previously essentially inaccessible link.

The platform’s graph-theoretic analysis revealed a small-world topology with densely connected hub complexes, indicating molecular-functional specialization and increased global communication capacity across scales^30,33,51^. Furthermore, by combining DPT and CAT analyses, MEA-seqX traced the pseudotemporal ordering along the developmental trajectory of cells within their spatial contexts and the progression of large-scale neural activity, providing unique insights into coordinated spatiotemporal dynamics in the hippocampal circuitry. MEA-seqX also demonstrated the potential for cell-type identification and highlighted the heterogeneity of neural cell types and their network-wide spectral fingerprints in the hippocampal circuit. The platform’s predictive capabilities using machine-learning algorithms allowed accurate forecasting of network-wide electrophysiological features based on spatial gene expression, showing a multiscale causal relationship between specific gene expressions and neural activity to offer a deeper understanding of neural dynamics, which could open new avenues for research in machine-learning and artificial intelligence. Combining MEA-seqX with intelligent neural networks may enhance our understanding of complex data, decision-making processes, and learning mechanisms^52,53^.

Furthermore, the potential of MEA-seqX in personalized medicine is promising. The identification of spatiotemporal transcriptional-electrophysiological biomarkers may aid in the early diagnosis and treatment of neurological disorders, ultimately leading to precision personalized therapies^54^. Moreover, exploring MEA-seqX in the context of other neurological and psychiatric disorders can deepen our understanding of disease mechanisms and therapeutic targets. The platform’s potential to uncover molecular-functional changes associated with various disease states may pave the way for developing novel treatments.

Looking ahead, several opportunities exist to enhance the capabilities of MEA-seqX. Continued advancements in technology and data analysis algorithms could further improve the platform’s resolution and predictive accuracy. Integrating additional modalities, such as proteomic and epigenomic data, could provide even more comprehensive insights into neural information processing^52^. The MEA-seqX platform’s adaptability goes beyond neural tissues, extending to electrogenic tissues like the olfactory bulb and cardiac tissues. This enables studying gene expression and electrophysiology interactions in diverse contexts. Its application in cardiac tissue offers insights into heart function, aiding cardiology advancements.

While the current implementation of MEA-seqX exploits the CMOS-MEA (4096 electrodes)^13,16^ built upon active-pixel sensor technology^55^, the platform’s scalability transcends reliance on a singular technology. It is adaptable to accommodate a spectrum of high-density technologies, such as those offered by switch-matrix technology (26,400 electrodes)^56^, among others. Importantly, MEA-seqX’s purview could extend beyond *ex vivo* applications, encompassing expansive *in vivo* investigations. Integration with state-of-the-art *in vivo* probes, such as Neuropixels^57^, SiNAPS^58^, or other emerging modalities, offers the potential to study functional neural dynamics and gene expression patterns within living organisms, bridging the gap between laboratory findings and real-world biological contexts.

## Materials and Methods

### Animals and Acute Brain Slice Preparation

All experiments were performed on 12-week-old C57BL/6J mice (Charles River Laboratories, Germany) in accordance with the applicable European and national regulations (Tierschutzgesetz) and were approved by the local authority (Landesdirektion Sachsen; 25-5131/476/14). Female C57BL/6J mice were obtained at five weeks of age and randomly distributed into two experimental groups – standard housed (SD) and enriched environment (ENR) housed as previously described^16^. ENR-housed mice lived in a specially designed cage containing rearrangeable toys, maze-like plastic tubes, tunnels, housing, and additional nesting material. This ENR cage environment has been shown to promote the enhancement of experience-dependent plasticity through increased stimuli and differentiated social interactions. Mice stayed in the assigned environment for six weeks before the experiments began and remained until their experimental date. Acute brain slices were prepared according to our previous studies^13,16^. Briefly, mice were anesthetized with isoflurane before decapitation. The brain was carefully removed from the skull and placed in a chilled cutting sucrose solution prior to slicing. The brain was placed into a custom-made agarose container and fixed onto the cutting plate. Dorsal, horizontal slices (300 μm thick) were prepared using Leica Vibratome VT1200S (Leica Microsystems, Germany). Slices were cut at 0–2 °C in an aCSF solution saturated with 95% O_2_ and 5% CO_2_ (pH = 7.2–7.4) of a high sucrose solution containing in mM: 250 Sucrose, 10 Glucose, 1.25 NaH2PO4, 24 NaHCO_3_, 2.5 KCl, 0.5 Ascorbic acid, 4 MgCl_2_, 1.2 MgSO_4_, 0.5 CaCl_2_. Next, HC slices were incubated for 45 min at 34 °C and then allowed to recover for at least 1 hour at room temperature before being used for network electrophysiology (n-Ephys) recordings with a high-density neurochip. A perfusate solution used during recordings contained in mM: 127 NaCl, 2.5 KCl, 1.25 NaH2PO_4_, 24 NaHCO3, 25 Glucose, 1.25 MgSO_4_, 2.5 CaCl_2_, and was aerated with 95% O_2_ and 5% CO_2_.

### Extracellular n-Ephys Recordings

All electrical recordings were performed using high-density CMOS-based biosensing MEA chips (CMOS-MEA) and an acquisition system (3Brain AG, Switzerland) customized to our recording setup. CMOS-MEA are composed of 4096 recording electrodes with a 42 μm pitch size to compose an active sensing area of ~7 mm^2^. The on-chip amplification circuit allows for 0.1–5 kHz band-pass filtering conferred by a global gain of 60 dB sufficient to record slow and fast oscillations^13^. For extracellular recordings, slices were moved and coupled onto the chip using a custom-made platinum harp placed above the tissue. To minimize experimental variation and maintain slice longevity, a heat-stabilized perfusion system delivered oxygenated recording perfusate to the neurochip with a flow rate of 4.5 mL/min and was temperature controlled at 37 °C throughout the experiment and recordings. We collected extracellular recordings at 14kHz/electrode sampling frequency and 1Hz recording frequency from spontaneous network-wide activity through pharmacological-induced evoked responses using 100 μM 4-aminopyridine (4AP)^16^ (Sigma-Aldrich, Germany). All solutions were freshly prepared, and pharmacological compounds were dissolved into the recording perfusate for the experiment. A custom-designed modular Stereomicroscope (Leica Microsystems, Germany) was incorporated into the system to capture the acute slice light imaging simultaneously with the whole HC circuit extracellular n-Ephys recordings. During offline analysis, these images were used to maintain the spatial organization of the brain slice tissue relative to the n-Ephys electrode layout.

### Spatially-resolved Transcriptomics in Hippocampal-cortical Slices

Spatially resolved HC sequencing and transcriptomic analysis were performed using SRT Visium gene expression slides (10X Genomics, USA). SRT slides are composed of 4 distinct capture areas containing 5,000 spatially barcoded spots with a 55 μm diameter to compose a capture area of ~6.5 mm x 6.5 mm sufficient for placement of the entire mouse HC slice. Immediately following n-Ephys recordings, slices were embedded in ~6.5 mm x 6.5 mm Tissue-TEK Cryomold containing Optimal Cutting Temperature (OCT) solution, frozen over dry ice, placed in a WHEATON CryoELITE tissue vial, and stored at −80 °C to maintain tissue stability and viability until SRT experimental date. Here, to optimize the number of cells per spot and provide a clear transcriptomic profile, slices were horizontally cryosectioned to 18 μm using Thermo Fisher Cryostar NX70 (Thermo Fisher Scientific, USA). Tissue was mounted on the SRT gene expression slide, and methanol was fixed at −20 °C for 30 minutes. Slices were hematoxylin and eosin (H&E) stained, and bright-field imaged to obtain morphological slice images. Following imaging, the slices were enzymatically permeabilized for 22 minutes on a thermocycler, and the resultant released mRNA was bound to the thousands of spatially barcoded mRNA-binding oligonucleotides within each spot. To generate cDNA from the oligonucleotide-bound mRNA, an enzymatic reverse transcription mixture (10X Genomics, USA) was applied and incubated in a thermocycler at 53 °C for 45 minutes. To generate cDNA second-strand synthesis, an enzymatic second-strand mixture (10X Genomics, USA) was applied and incubated in a thermocycler at 65 °C for 15 minutes. To denature enzymes, a basic elution buffer (EB) (Qiagen, Germany) with a pH of 8.7 was applied, and the final sample was stored in a corresponding tube per capture area containing Tris-HCl. Finally, the spatially barcoded, full-length cDNA is prepared for library sequencing through PCR amplification. To determine the optimal cycle number (Cq) via qPCR, a qPCR mix using KAPA SYBR FAST (Kapa Biosystems, USA) and a 1 μL sample from each cDNA sample was added to a clean qPCR plate. Following the incubation protocol, a Cq value of 15.7 was determined for the cDNA sample, which corresponded to 16 amplification cycles. An amplification mixture (10X Genomics, USA) was added to the cDNA sample tubes, and the qPCR amplification protocol was completed according to the obtained Cq value. Samples were stored overnight at 4 °C overnight before proceeding to sequencing. Library construction and sequencing were carried out at the Dresden Concept Genome Center (DcGC) using the HiSeq 2000 Next Generation Sequencer (Illumina, Inc., USA). Sequenced data was processed with the Space Ranger (10X Genomics, USA) pipeline to recreate the spatial arrangement, which aligns the H&E stained bright-field image with the spatially barcoded gene expression data based on the fiducial spots in the slide capture area border. The pipeline performs alignment, tissue detection, fiducial detection, and barcode/UMI counting.

### Data Analysis

All basic and advanced algorithms used in this work were developed and implemented with custom-written Python scripts. Any package add-ons are cited accordingly.

### SRT Quality Control and Gene Expression Normalization

Prior to data analysis, technical batch effects and experimental variation were ruled out using a single-cell analysis toolkit, Seurat^59^, and an add-on package STutility (https://ludvigla.github.io/STUtility_web_site/). These packages statistically quantified the number of unique genes (nFeature RNA) and the number of UMIs (nCount RNA) across all samples and conditions. To further delineate and find shared hippocampal structures between the two conditions, a further add-on package, Harmony, recomputed the UMAP embedding and clustering to return an integrated low-dimensional representation of the data^60^. As each dataset was found to have ~5500 median genes per spot, SRT spots in each dataset with fewer than 1000 unique genes were filtered out of the analysis. Next, to downsize the total number of genes for analysis, mitochondrial and ribosomal protein-coding genes were filtered out of analysis. Finally, to account for technical batch variation and detect highly variable genes, overall gene expression per SRT spot was normalized by total counts of each gene over all SRT spots so that each spot has the same count after normalization. This was implemented using the scanpy.pp.normalize_total python package and is available on GitHub (https://github.com/theislab/scanpy)^61^.

### Oscillatory Pattern Detection and Waveform Classification

Prior to data analysis, oscillatory patterns of LFPs were detected in each recording with hard threshold algorithms^16^. Furthermore, detected events were further processed and filtered with a low-pass 4th-order Butterworth filter (1-100Hz). Finally, quantile thresholding was employed in a custom-written Python script to remove spuriously firing electrodes or non-physiologically detected events^16^. To characterize and allocate the distinct features and shapes of the recorded LFP oscillatory waveforms to specific interconnected HC layers, we implemented Principal Component Analysis (PCA) and K-means clustering algorithms in a procedure as previously described^13^.

### Structural Clusters

To characterize local and global hippocampal subnetwork behavior, the functional firing n-Ephys electrodes were structurally related to a specific HC region through an overlay of light microscope hippocampal images on the CMOS-MEA layout. Electrodes were then grouped into clusters based on structural markers on the HC slice – DG, Hilus, CA1, CA3, EC, and PC^16^. To characterize the transcriptomic profile in these six major regions, SRT spots were structurally related to a specific HC region through an overlay of H&E-stained bright field microscope images on SRT spot layout using Loupe Browser (10X Genomics, USA).

### Multiscale Spatial Alignment

To infer a correspondence between the n-Ephys electrode-SRT spot interface and their respective network-wide functional electrical activity and transcriptomic feature readouts with spatial localization, MEA-seqX implements a multiscale spatial alignment procedure. To provide the transcriptomic and electrophysiologic profiles of the same network with spatial context, automatic slice alignment is performed using image resizing and rotation. This alignment is based on optical imaging, n-Ephys electrode-SRT spot interface physical size, and related hippocampal-cortical structural inputs to put the multiscale data in the same dimension. First, MEA-seqX implements an automatic scaling algorithm based on n-Ephys electrode-SRT spot sizes to resize the light H&E-stained bright field microscope slice image from SRT to the respective light microscope hippocampal image from n-Ephys. Importantly, the n-Ephys electrode-SRT spot matching is not one-to-one due to the difference in technology resolution; instead, it is a fractional matching based on related hippocampal-cortical structural inputs. As such, each SRT spot has related n-Ephys electrodes with averaged electrophysiological features from the related electrodes. Next, slice spatial alignment and rotation are computed between the two slice images with the following procedure – i) Hippocampal-cortical structural reference points {*i*, *j*} and {*k*, *l*} are assigned for each scale, SRT and n-Ephys respectively, where {*i*, *k*} are midpoints in the DG crest and {*j*, *l*} are on the DG supra blade edges. ii) SRT reference point *i* is aligned with n-Ephys reference point *k* to place both scales in one dimension. ii) Following the alignment of reference points {*i*, *k*}, a final alignment for reference points {*j*, *l*} is based on the difference between *θ_n−Ephys_* and *θ_SRT_*. iv) Given that the coordinates of both arrays are known, the distance of *x*, *x*′, *y*, and *y*′ is used to calculate the angle of *θ_SRT_* and *θ_n−Ephys_*. v) To determine *θ_SRT_*, the horizontal intersection line between aligned reference points {*i*, *k*} was used to define *x* while a vertical intersection line between reference point *j* and the horizontal intersection line was used to define *y*. vi) To determine *θ_n−Ephys_*, the horizontal intersection line between aligned reference points {*i*, *k*} was used to define *x*′ while a vertical intersection line between reference point *l* and the horizontal intersection line was used to define *y*′. vii) The final angle of rotation is defined as *θ_iNeuromics_* = *θ_n−Ephys_* − *θ_SRT_*, which, when applied, align reference points {*i*, *k*} as the final multiscale reference point *m* and align reference points {*j*, *l*} as the final multiscale reference point *n*.

### Mean Activity n-Ephys Features

To determine how spatial gene expression patterns are related to a functional n-Ephys feature, filtered genes were correlated with one of the network features using Spearman’s correlation^62^ and sorted according to significance using Benjamini-Hochberg false discovery rate adjusted p-value^63^. Functional network features of large-scale spatiotemporal LFP oscillations included LFP rate, amplitude, energy, delay, and positive and negative peak count^13,16^.

### Targeted Gene Lists

Specific gene lists were formulated based on functional gene ontologies. Genes related to families of immediate early genes, signaling pathways, hippocampal function, and neurogenesis were compiled into six lists^24,25^.

### Non-Negative Matrix Factorization (NMF)

We implemented an unsupervised machine-learning algorithm using a sparsity-constrained non-negative matrix factorization to identify individual spatiotemporal patterns emerging from SRT and n-Ephys networks^26^. NMF factorization is described and adapted from the Scikit-learn 1.2.2 python package (sklearn.decomposition.NMF)^64^. First, the input V-matrix contains the combined information from SRT (i.e., gene expression values from the IEG gene family) and n-Ephys (i.e., LFP rate or Degree network feature), with each data entry comprising the expression value of each gene related to network feature value (n) with spatial localization (m).

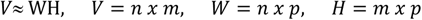

Where *n* is the spatially localized spots, *m* is the related gene expression to network feature, and *p* is the number of factors.

The resultant decomposed basis W-matrix contains the spatial gene expression patterns and their locations related to the functional feature, and the coefficient H-matrix represents the contribution of these basis vectors to each spatial location and n-Ephys feature. To optimize the distance between *V* and the product matrices *H* and *W*, we implemented the widely used distance optimizing function squared Frobenius norm (F), which added sparsity constraints for the factors^64^.

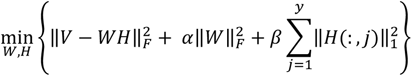

Where *W_ij_* and *H_ij_* are nonnegative value (1 ≤ i ≤ x, 1≤ j ≤ y). α and β are the corresponding regularization parameters for *H* and *W*.

### Mutual Information

To present the collectivity of spots in a multilayered network based on gene expression, gene expression distribution was calculated for each target gene list. Next, mutual information distance scores were calculated for the gene information from each target gene list in each spot, compared between spots, and sorted by cluster^28^. We computed the mutual information using adapted functions from Scikit-learn Scikit-learn 1.2.2 python package (sklearn.metrics.normalized_mutual_info_score)^64^.

### Functional Connectivity

To infer large-scale statistically dependent connectivity for functional firing activity over a multilayered hippocampal network, cross-covariance was calculated between pairs of firing n-Ephys electrodes using Pearson’s correlation coefficient (PCC) followed by Directed Transfer Function (DTF) and Multivariate Granger causality as previously described^13,16^.

### Graph Map Visualization

To visualize large-scale network connectivity in both SRT and n-Ephys datasets, the mutual information distance scores and functional connectivity data architecture, respectively, were constructed to contain nodes and edges as described previously^16^. The data were converted into (.gexf) file format and were directly read and visualized in the Gephi program 9.2 version (https://gephi.org). To study the functional interactions of selected genes over the spatial network array, we filtered the mutual information scores between all paired spots to include those over the mean and two standard deviations. Therefore, the threshold value was set to include genes with r ≥ 0.8 for each target gene list. To examine the functional connections of n-Ephys electrodes, the top 2% of the total functional links were included. Both SRT and n-Ephys connectivity maps were plotted with similar edge weights and degree range queries.

### Network Topological Metrics

Graph Theory was used to characterize overall network topology and interconnectedness based on functional connectivity from n-Ephys detected LFP events or mutual information scores from SRT gene expression. We computed the topological metrics in custom-written Python code, as we previously reported^13,16^. Briefly, the network connectivity topological metrics are described by considering the node *n* as the central component of the graph that may or may not be connected to one another. In our case, a node *n* corresponds to a specific n-Ephys electrode or SRT capture spot in the sensing arrays, where the edges *e* are the functional links or connections between each node *n.* To present overall network topology and features, we selected the following graph theory topological parameters:

#### Degree

To characterize the different representations of network connectivity, we characterized the degree *k* of a node *n* to describe the number of edges connected to a node as previously described^16^.

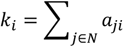

Where *k_i_* denotes the degree of a node *i*. *a_ij_* denotes the connection between nodes *i* and *j*. *N* is the set of all computed nodes in the network.

#### Hub Nodes and Rich Club Nodes

To determine centralized, important nodes in a network and reveal network topology, hub nodes and rich club nodes were analyzed. Hub nodes were detected based on three nodal metrics - node strength, clustering coefficient, and network efficiency. The metric value for each node was calculated and compared to determine whether the node value was in the top 20% of all nodes^16^. To restrict the definition of the hub node, we set limits with a hub score. Our hub score was valued between 0 and 3, where nodes either satisfied the top 20% in none, 1, 2, or all 3 nodal metrics. Within the hub node group is a subgroup of nodes with dense connections that conferred the rich-club nodes and are described as hub nodes with a higher degree than the average and provided by the rich club coefficient *ϕ*(*k*)^16^

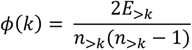

Where *k* denotes the degree, *n*_>*k*_ represents the number of nodes whose degree is larger than a given value *k*, and *E*_>*k*_denotes the number of connections in a subnetwork comprising *n*_>*k*_.

### Network Topology Characterization

To determine the potential impact of hub nodes on the network function and the organizational processes shaping network topology, we characterized the degree distributions *P(k)* of detected nodes in n-Ephys and SRT datasets, which resulted in decayed distribution with a power-law tail^65^.

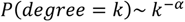

To estimate the power-low degree distribution *P(k)* to describe the scale-free topology with a small-world attribute, we used the lognormal model fit.

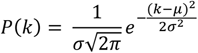

Where *μ* and *σ* are the mean and standard deviation of the distribution, respectively. To visualize the best-fit network characterization, a complementary cumulative distribution function (cCDF) was used instead of the probability density of the node degree and plotted on logarithmic axes for a more robust visualization of the high-k regime. Goodness-of-fit tests were performed between actual data and fitted models and were estimated by the coefficient of determination *R*^2^. Finally, Pareto linear binning (scipy.stats.pareto)^66^ was applied to discretize the power law distribution.

### Diffusion Pseudotime

To pinpoint dynamic transcriptional changes from static, spatially resolved sequencing data and to determine the impact of intrinsic and extrinsic influences on the distinct dynamic process under examination, we employed diffusion pseudotime (DPT)^36^. DPT uncovers the underlying dynamics of biological processes and, in this case, the temporal trajectories of specific gene expression from spatially resolved hippocampal transcriptomes. While traditional diffusion maps effectively denoise data while maintaining the local and global structure, the resultant maps usually encode the information in higher dimensions, limiting the visualization. To overcome this prior to the employment of DPT analysis, we implemented the potential of heat diffusion for affinity-based transition embedding (PHATE) on spatial transcriptomic data, which presents information at lower dimensionality (https://github.com/KrishnaswamyLab/PHATE)^67^. PHATE encodes both local data and global data in a manifold structure. Local data relationship similarities are encoded by applying a kernel function on Euclidean distances. Global data relationships are encoded via potential distances where the local similarities are transformed into probabilities. These diffusion probabilities are determined by transforming the local information into the probability of transitioning from one data point to another in a single step of a random walk. This can be powered to t-steps to give t-step walk probabilities for both local and global distances. In our dataset, each spatial spot has a determined relationship to each nearest neighbor or distant spot in a weighted graph^67^. DPT analysis then orders the transcriptomic spots according to the probability of differentiation toward a different spot^36^.

### Cell-Type Deconvolution

To determine cell type colocalization and examine differences between two hippocampal transcriptomes, conditional autoregressive-based deconvolution (CARD) was performed using a single-cell sequencing reference^40,41^. The CARD-based analysis is found on GitHub (https://github.com/YingMa0107/CARD) and was adapted for spatial transcriptomic data in Python. Within the reference was a broad group classification of hippocampal cell types, including astrocytes, endothelial cells, ependymal cells, macrophages, microglial cells, neurogenic cells, neurons, oligodendrocytes, and polydendrocytes with the accompanying subgroups. Low-count cell type filtering downsized 85 to 76 hippocampal cell types.

### Prediction with XGBoost Algorithm

To determine if specific gene expression values can predict related HC network feature parameters per SRT spot, such as LFP rate, amplitude, and duration, we implemented the Gradient Boosting (XGBoost) Algorithm, which integrates multiple tree models and has a strong interpretability^48^. XGBoost is described in the Scikit-learn 1.2.2 Python package (sklearn.ensemble.GradientBoostingClassifier) with training and testing datasets implemented as described in the package (sklearn.model_selection.train_test_split)^64^. These datasets contained the spatially resolved gene expression values based on specific gene lists and the related network features from functional n-Ephys datasets. To level input information between the SD and ENR datasets for better comparison, half of the ENR dataset was randomly generated to use as input for the training and testing datasets. This normalization was selected as the ENR has a two-fold difference in network feature parameters, as previously described^16^. The input datasets were split equally into an initial training dataset and a testing dataset, real data points trained over 100 iterations. The final predicted and real data points were used to determine the predictability, accuracy, and significance of network feature prediction from transcriptomic data^68^. Packages implemented for statistics included Scikit-learn 1.2.2 Python packages to calculate the prediction accuracy (sklearn.metrics.explained_variance_score)^64^ and Scipy 1.10.1 to calculate the Pearson correlation coefficient (scipy.stats.pearsonr) and (scipy.stats.ttest_ind)^66^. Accuracy results were compared over multiple final data outputs and values over the mean, and three standard deviations were determined to be within the threshold of chance.

### Statistical Analysis

All statistical analysis was performed with Originlab 2020 or as described in package add-ons. Data in this work were expressed as the mean ± standard error of the mean (SEM) unless otherwise denoted as standard deviation. Box charts are determined by the 25^th^-75^th^ percentiles and the whiskers by the 5^th^-95^th^ percentiles with lengths within the Interquartile range (1.5 IQR). Also, the lines depict the median and the squares for the mean values. Differences between groups were examined for statistical significance, where appropriate, using the Kolmogorov-Smirnov test, one-way analysis of variance (ANOVA), or two-way ANOVA followed by Tukey’s posthoc testing. P < 0.05 was considered significant.

## Authors Contributions

B.A.E. performed experiments, analyzed multiscale data, and generated the figures. X.H. wrote the code and analyzed multiscale data. D.K. performed SRT experiments. S.K. performed experiments. L.L. and J.L. helped analyze SRT data and discussed the results of the study. I.D. and J.F. helped with gene ontology and SRT analysis and discussed the results of the study. G.K. provided conceptual input, designed and performed enrichment experiments, supported interpretation of results, and contributed to co-funding. H.A. conceptualized, planned, and supervised the project, designed and performed experiments, developed computational tools, undertook formal multiscale analysis, and generated the figures. G.K., and H.A., jointly coordinated the study and wrote the manuscript. All authors reviewed and approved the final manuscript.

## Acknowledgments

This study was financed from basic institutional funds (DZNE) and partly from the Helmholtz Validation Fund (HVF-012). We would also like to acknowledge the platform for behavioral animal testing at the DZNE-Dresden (Alexander Garthe, Anne Karasinsky, and Sandra Günther) for their support.

