## Supplementary Data for "MEA-seqX: High-resolution Profiling of Large-scale Electrophysiological and Transcriptional Network Dynamics"

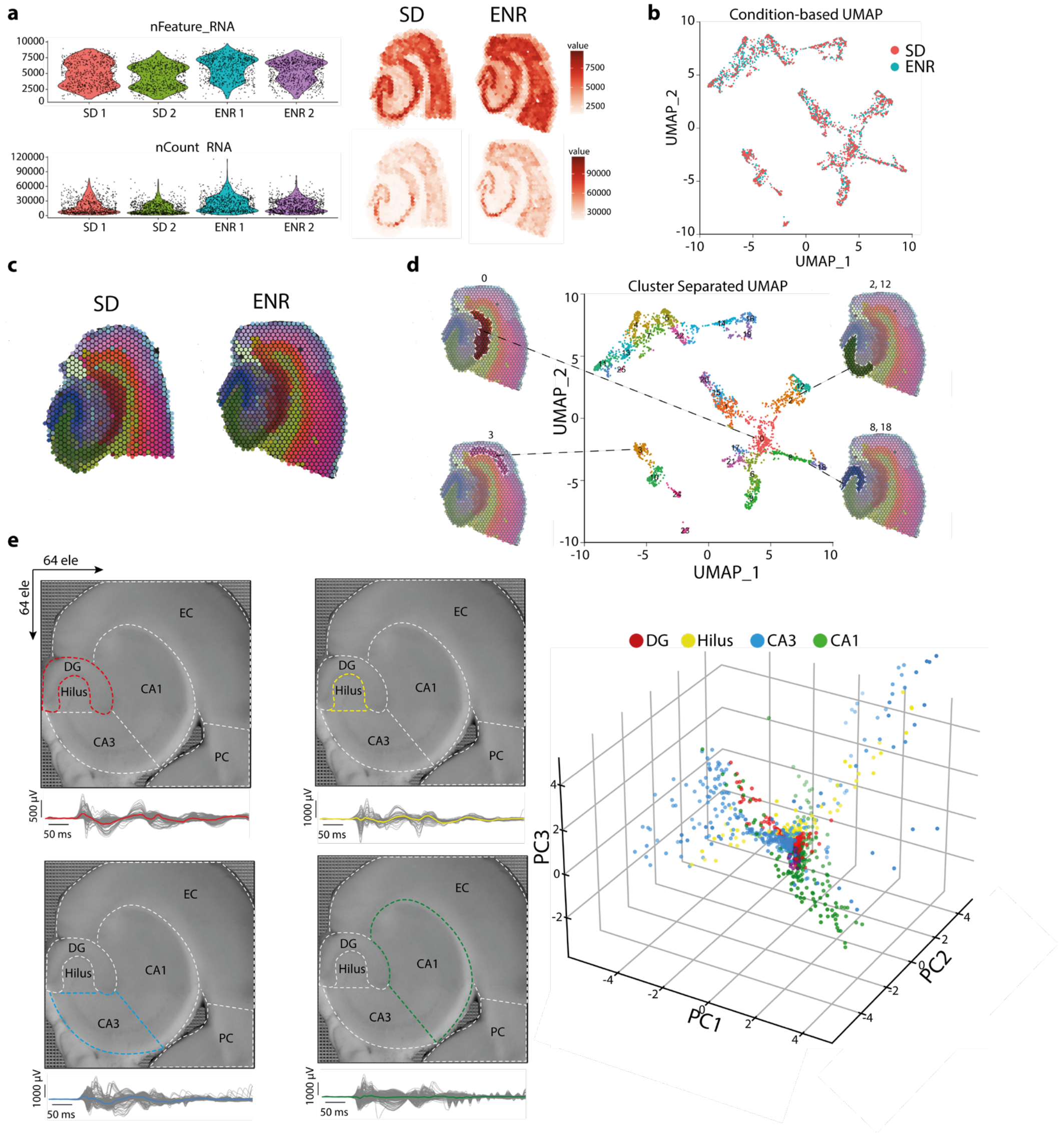


**Supplementary Figure 1. a)** The nFeature RNA and nCount RNA Statistics, along with feature maps, present the number of distinct genes and the number of unique molecular identifiers (UMIs) in SD (n=2) and ENR (n=2) transcriptomes. These distributions, part of quality control, highlight consistent counts of unique genes and UMIs across all samples and conditions, effectively mitigating potential technical batch effects and experimental variability. **b)** Employing both Harmony and Seurat programs, a condition-based UMAP analysis differentiates transcriptomic profiles based on conditions. The resultant UMAP reveals a congruent tissue structure between the two conditions, exhibiting precise cluster separation at high resolution. **c)** Segmentation of HC clusters based on spatial gene profiles is performed for SD and ENR brain slices. **d)** Selected clusters, aligned with DG, CA3, CA1, and EC regions, are positioned within a UMAP representation. This lower-dimensional depiction effectively showcases the separation of clusters based on gene expression profiles, capturing tissue heterogeneity and gradual expression variations. **e)** Through PCA analysis and the K-means algorithm, a classification of multilayered HC oscillatory waveform shapes is established. This classification is rooted in features extracted from LFP events, unraveling intricate insights into the diversity of waveform patterns across different layers of the hippocampus.

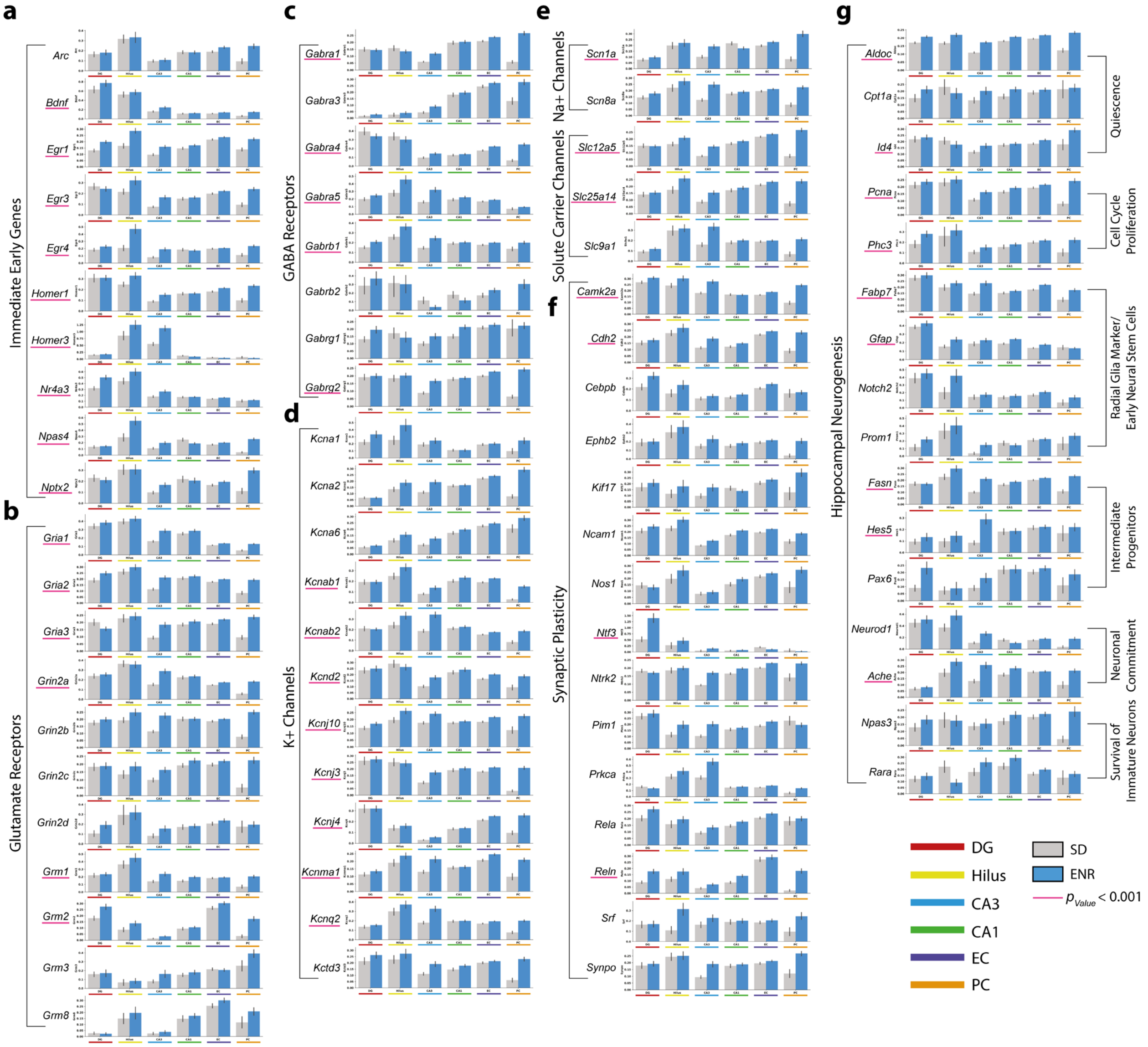


**Supplementary Figure 2. a-f)** Illustrate the elevated transcriptional expression observed in ENR compared to SD conditions across HC clusters. Genes from six distinct functional families, encompassing Immediate Early Genes (IEGs), Hippocampal Neurogenesis, Signaling Pathways, Receptors and Channels, Synaptic Plasticity, and Synaptic Vesicles, are systematically categorized. Notably, key genes critical for neuronal activity and function, such as *Bdnf*, *Egr1*, *Homer1*, *Npas4*, *Gria2*, and *Campk2a*, display heightened expression levels within the DG transcriptome of ENR, underscoring the enriched transcriptional landscape induced by environmental enrichment (*p < 0.001 ANOVA*).

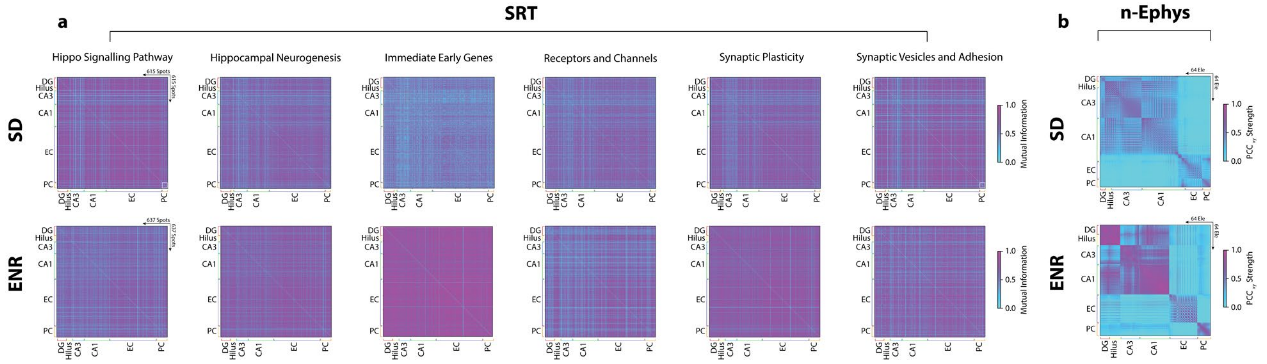


**Supplementary Figure 3. a)** Analysis of gene expression interplay matrices within six gene families using mutual information reveals an intriguing contrast between SD and ENR conditions. ENR displays heightened statistical transcriptional connectivity and coordinated activity compared to SD, suggesting a robust influence on HC-subnetwork relationships. **b)** Cross-covariance of pairwise firing electrodes is evaluated through PCC, unveiling intricate connectivity patterns. Compared to SD, ENR connectivity matrix exhibits amplified local and global spatiotemporal interactions, which harmonize with the cross-correlogram observations. These findings collectively shed light on the nuanced impact of ENR on the dynamic transcriptional-functional interplay within hippocampal subnetworks.

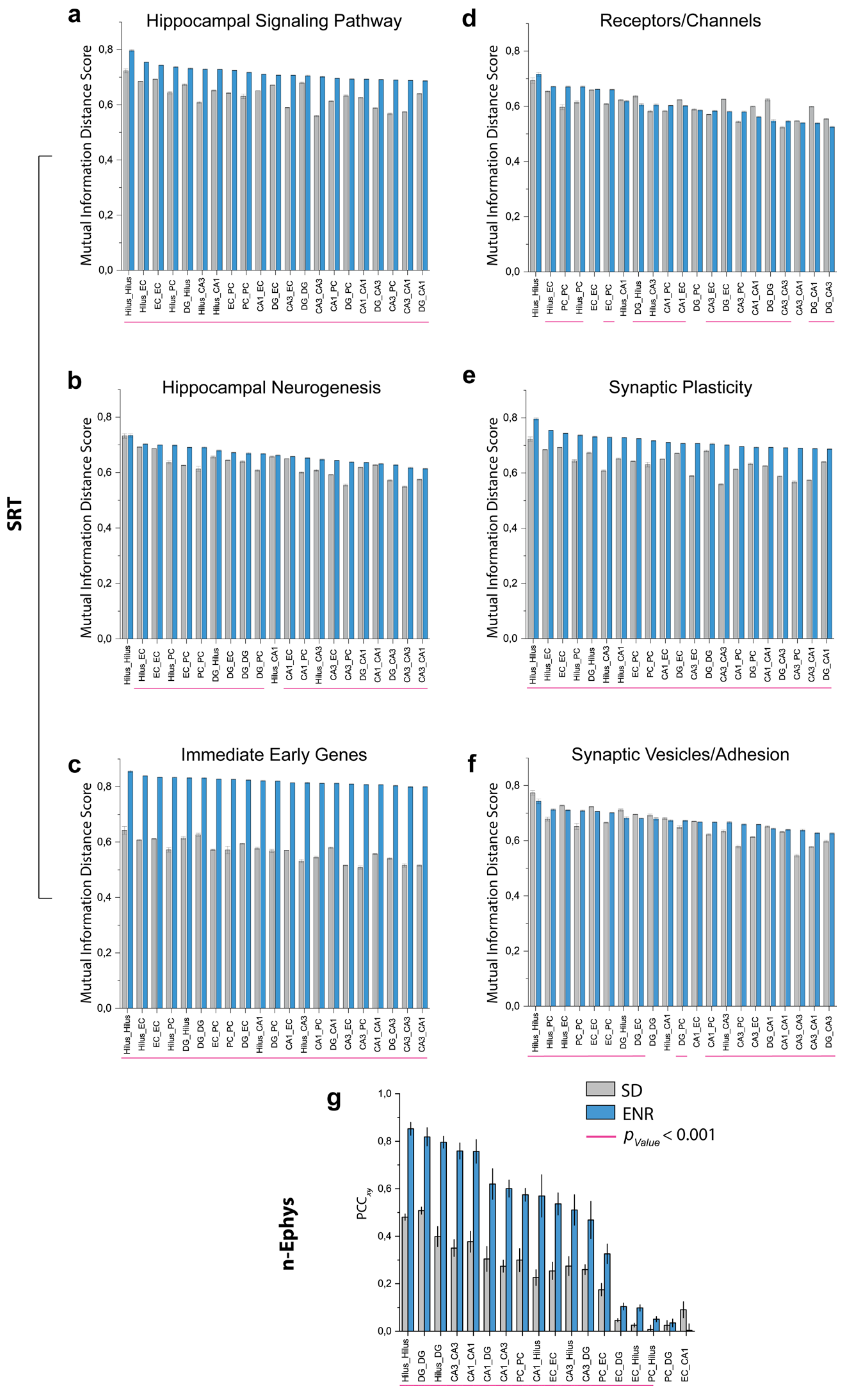


**Supplementary Figure 4. a)** Mutual information distance scores were computed for target gene families within SRT spots, revealing differences in interactions within HC subnetworks. The ENR transcriptome exhibits higher mutual information, indicating robust, coordinated activity and communication among gene expression patterns compared to SD (*$p < 0.001$ ANOVA*). **(b)** Correlation matrices are quantified using PCC of LFP activity across interconnected HC layers (*$p < 0.001$ ANOVA*). This corresponds with increased spatial interactions in ENR compared to SD networks, as shown in (a).

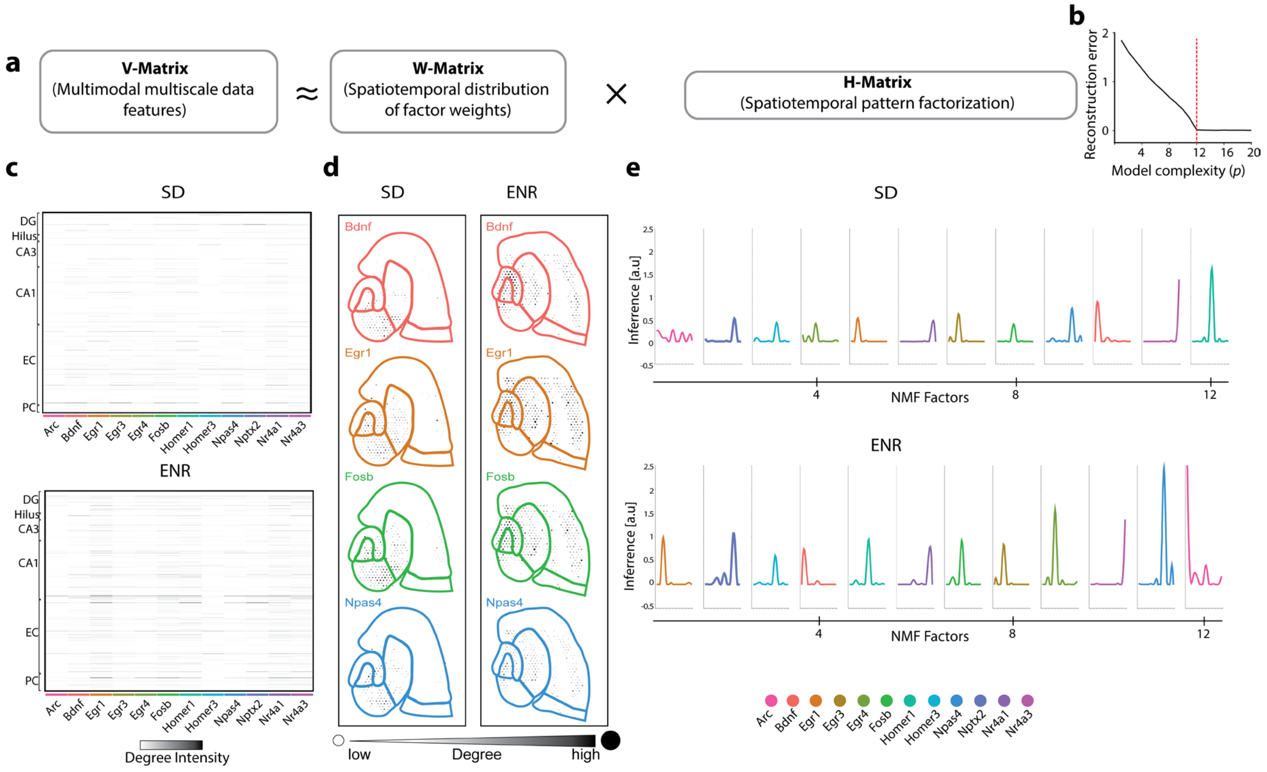


**Supplementary Figure 5. a)** The application of sparsity-constrained NMF analysis, as demonstrated in Figure 2c-f. **b)** The reconstruction error of V, considering various quantities of shared spatiotemporal patterns in H. An evident "elbow" is observed at n = 12, where the enhancement of the model diminishes. **c-e)** The analysis quantifies decomposed sets of genes, spatial locations, and connectivity node degree features, revealing simultaneous variations in network topological metrics linked to spatially-resolved IEG expression and network connectivity.

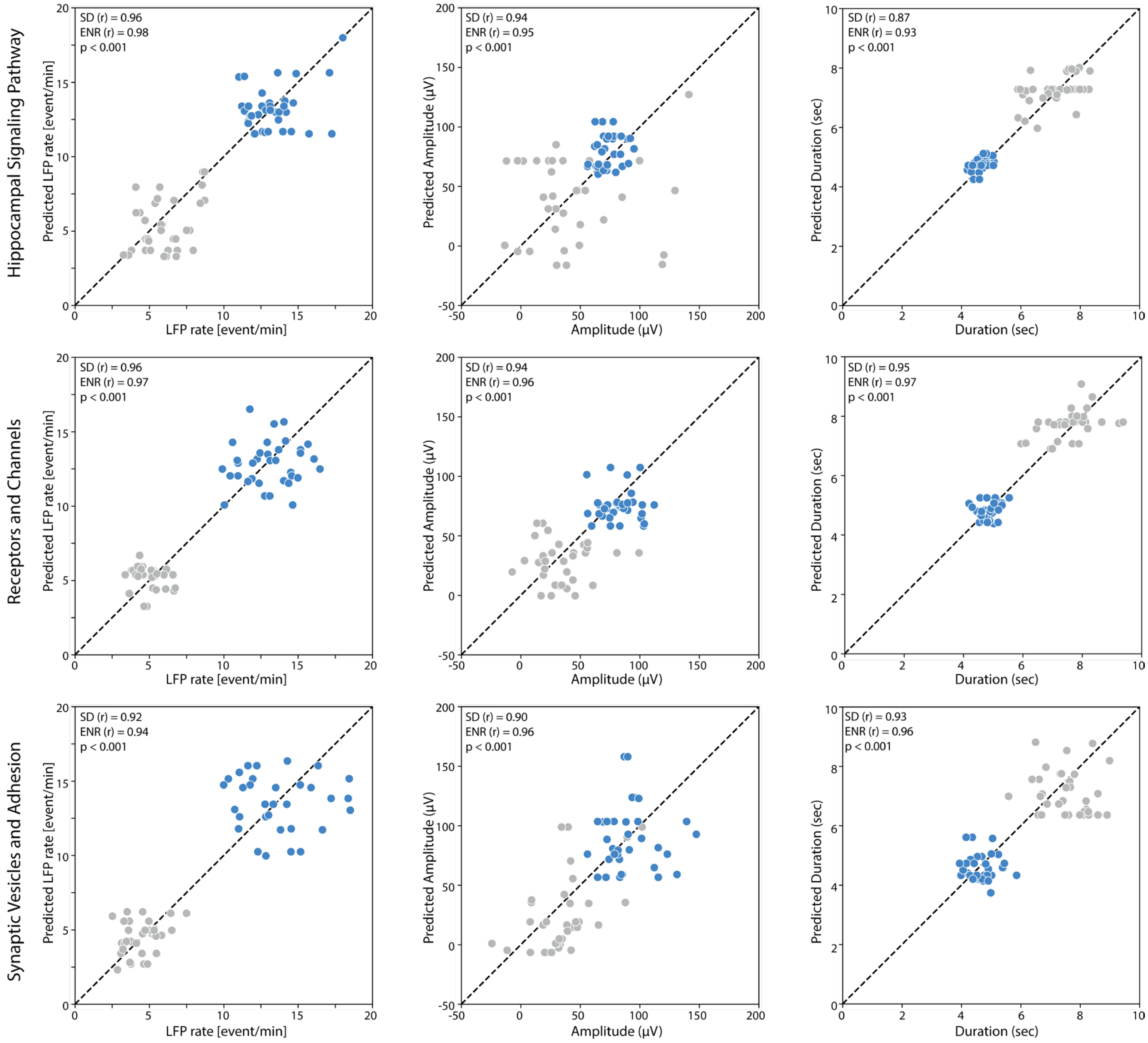


**Supplementary Figure 6.** Application of XGBoost algorithm to predict network electrophysiological metrics (LFP rate, amplitude, and duration) from transcriptomic data of each specified gene family from SD and ENR data. The prediction of n-Ephys metrics from the transcriptomics data is evaluated with Pearson correlation coefficient (*r*) in SD and ENR. The significant difference between the predicted SD and ENR values in all gene families is indicated ($p < 0.001$, *ANOVA*).
